## supplemental figure and table for "Elucidating ATP’s Role as Solubilizer of Biomolecular Aggregate"

### Supporting Information for “Elucidating ATP’s Role as Solubilizer of Biomolecular Aggregate”

**[1]Tata Institute of Fundamental Research Hyderabad, Telangana, India, 500046**

**[2]Indian Institute of Science Education and Research, Kolkata, Campus Rd, Mohanpur, Haringhata Farm, West Bengal 741246**

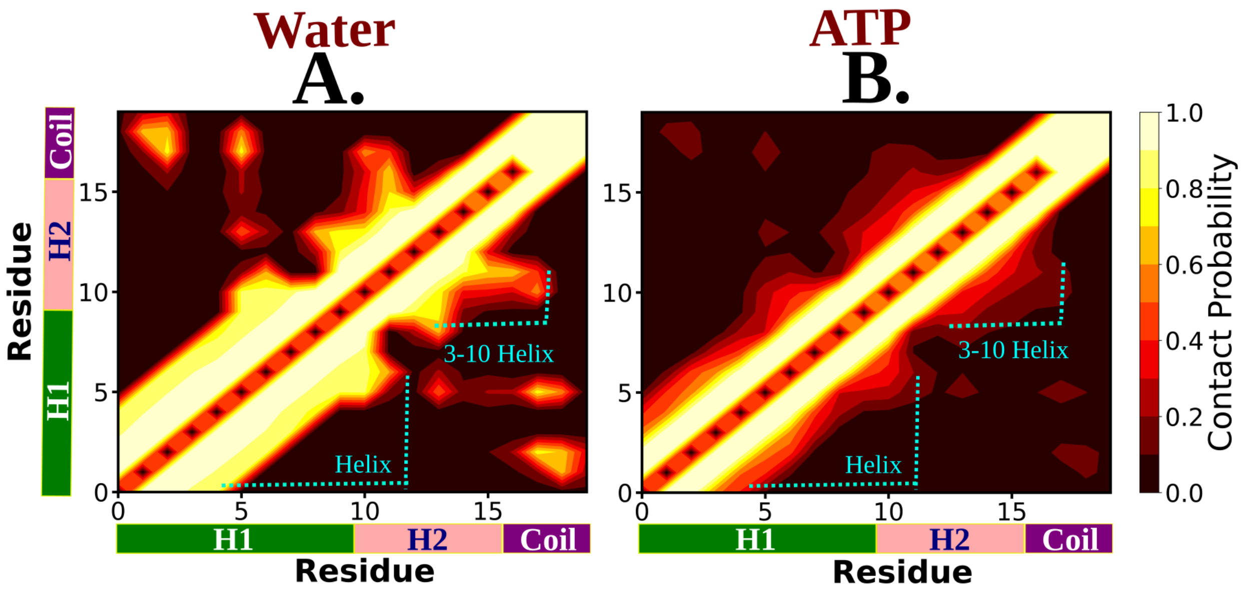


Figure S1: The residue wise contact map of Trp-cage monomer is shown for neat water and 0.5 M aqueous ATP solution in figure A and B respectively.

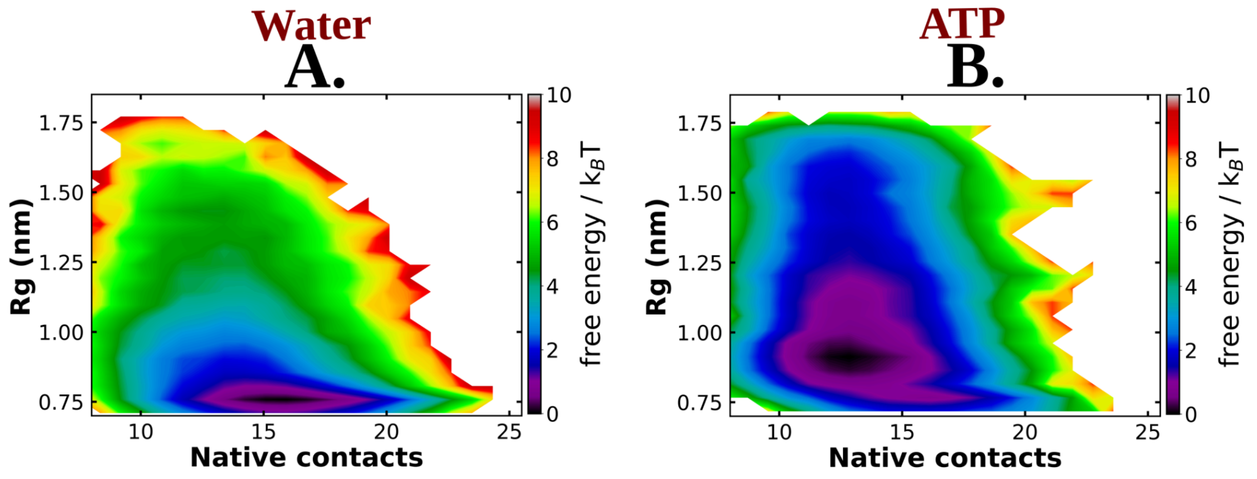


Figure S2: The 2D free energy profile of Trp-cage monomer estimated with respect to $R_g$ and native contacts are shown in figure A and B for Trp-cage in neat water and 0.5 M ATP respectively for the simulations with charmm36 force field.

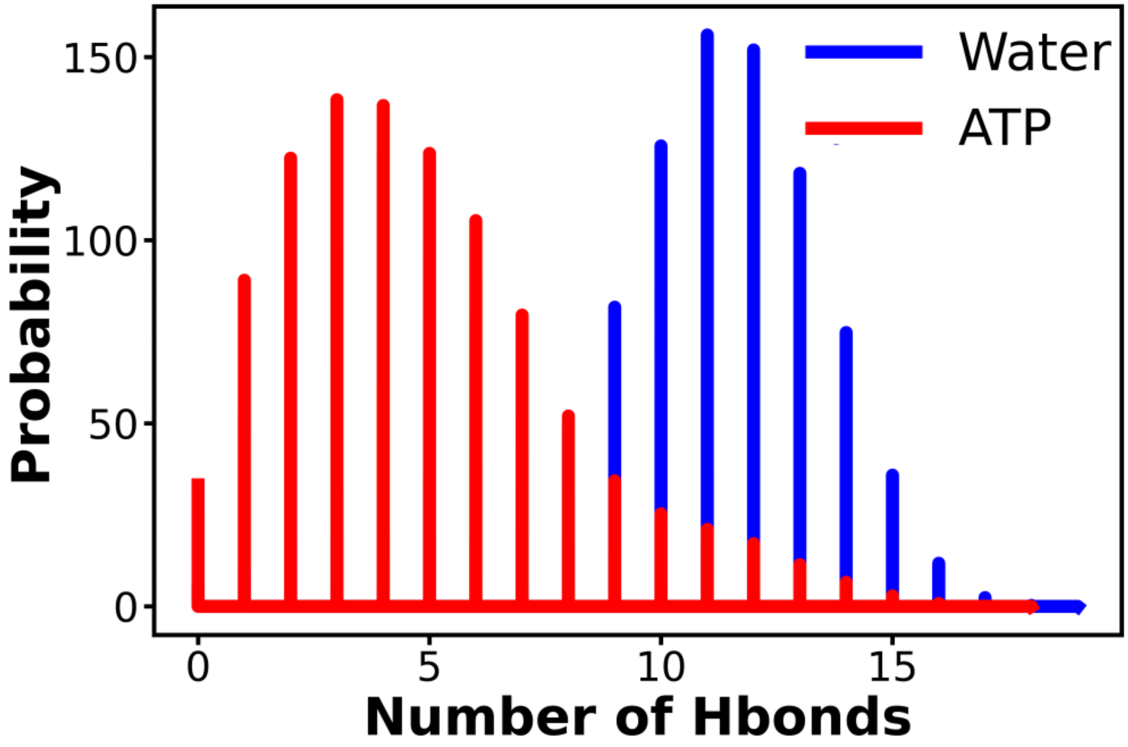


Figure S3: The probability distribution of the number of intrachain hydrogen bonds has been compared for neat water and in 0.5 M ATP solution.

### REMD+adaptive sampling simulation derived

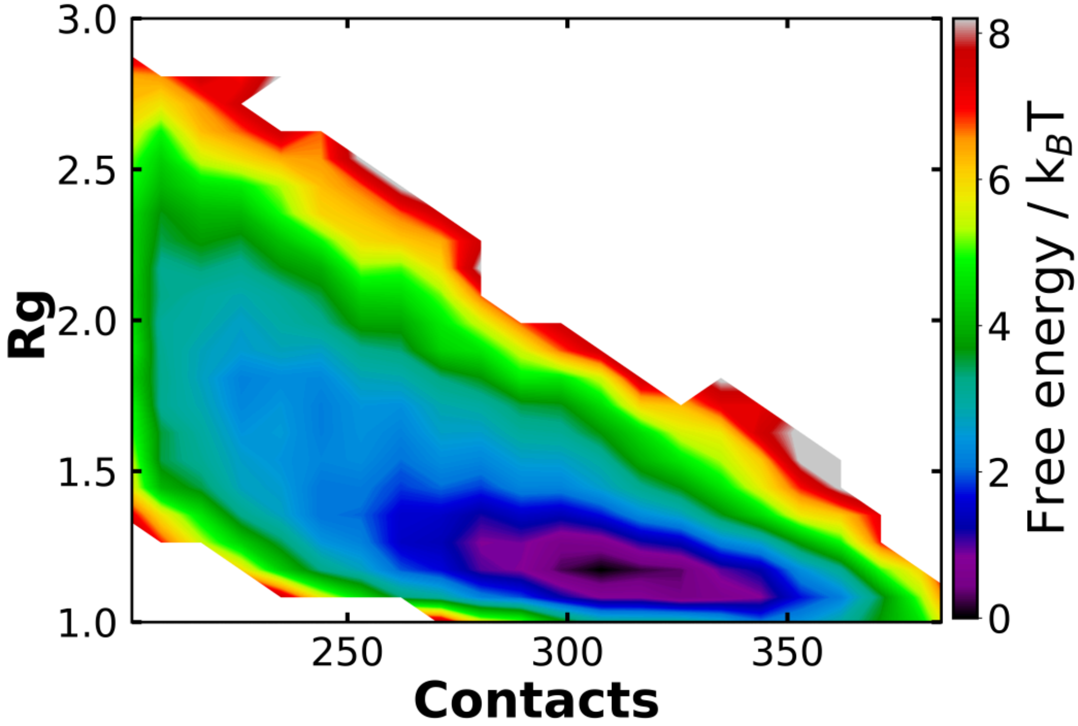


Figure S4: The 2D free energy profile estimated for Aβ40 protein in absence of ATP obtained from REMD simulation followed by adaptive sampling simulations.

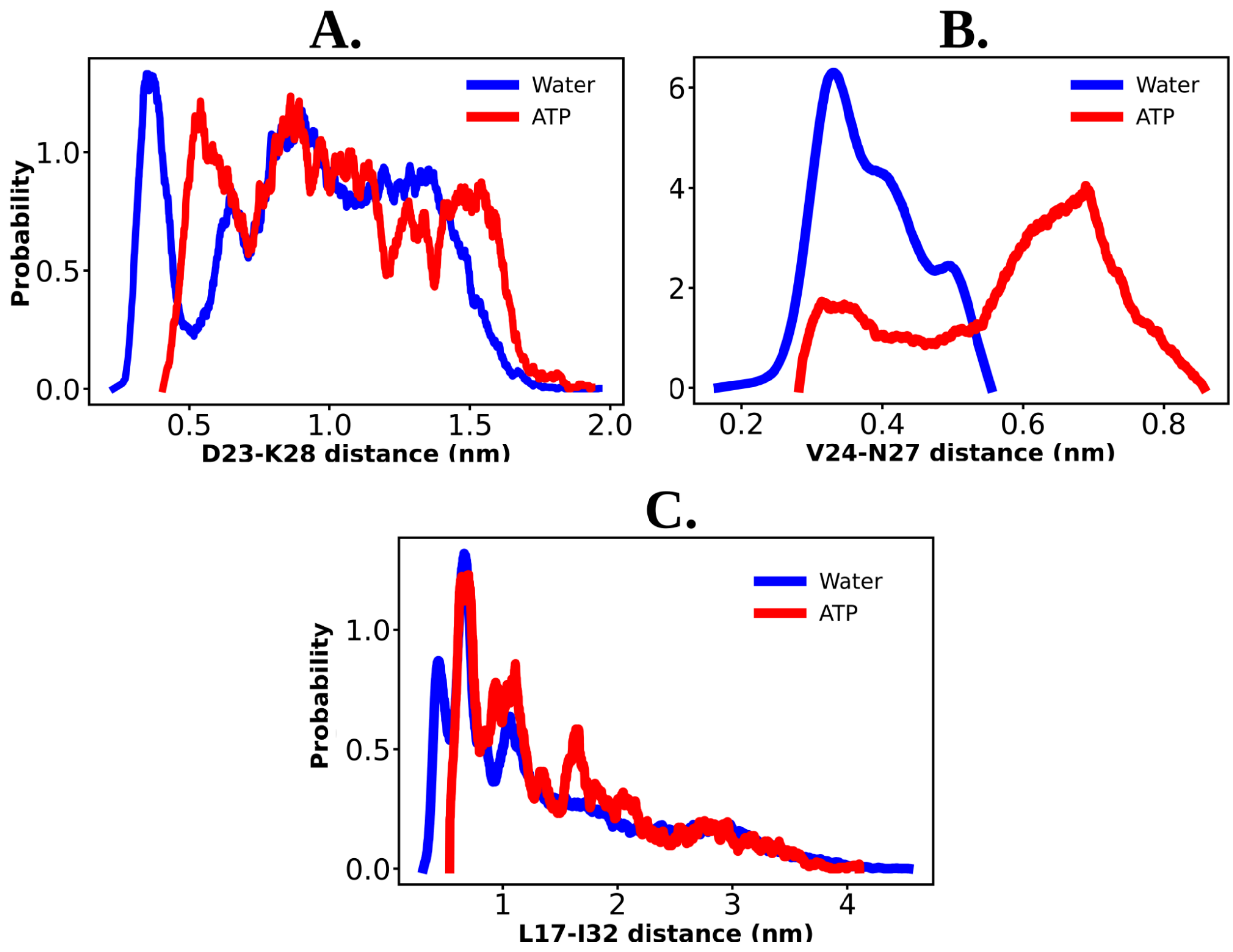


Figure S5: The probability distribution of the distance between the residue pair of D23-K28, V24-N27 and L17-I32 are shown for the protein Aβ40 both in 50 mM NaCl salt solution and 0.5 M aqueous ATP solution containing 50 mM NaCl salt in figure A, B and C respectively.

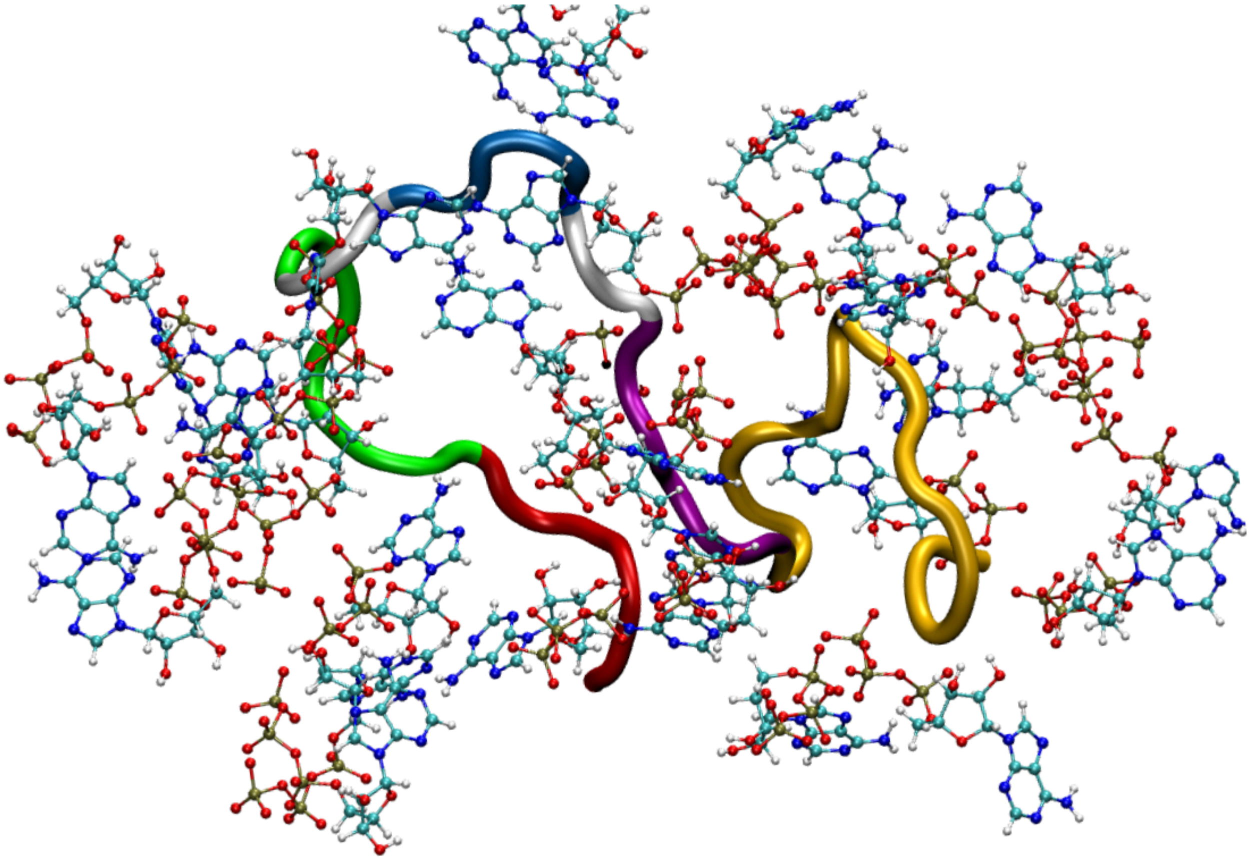

Figure S6: Figure shows a simulation snapshot representing ATP’s interaction with the Aβ40 protein.

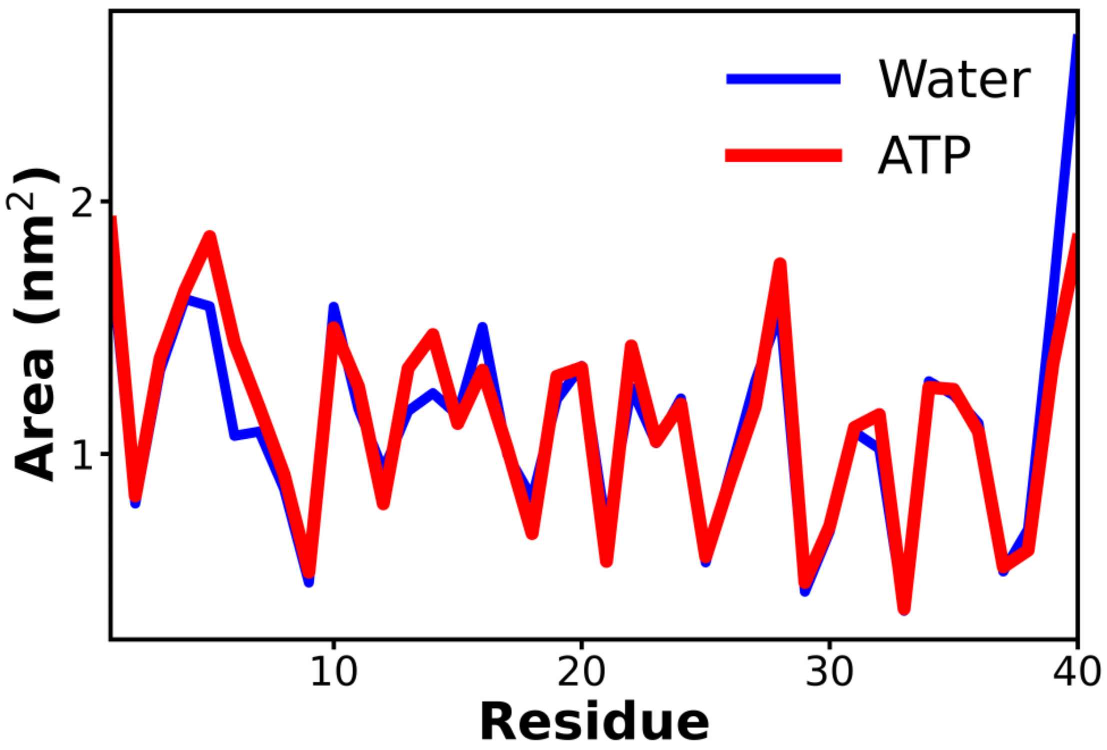


Figure S7: The comparison of the solvent accessible surface area of the protein Aβ40 has been shown for 50 mM NaCl salt solution (blue) and 0.5 M ATP in 50 mM NaCl salt (red).

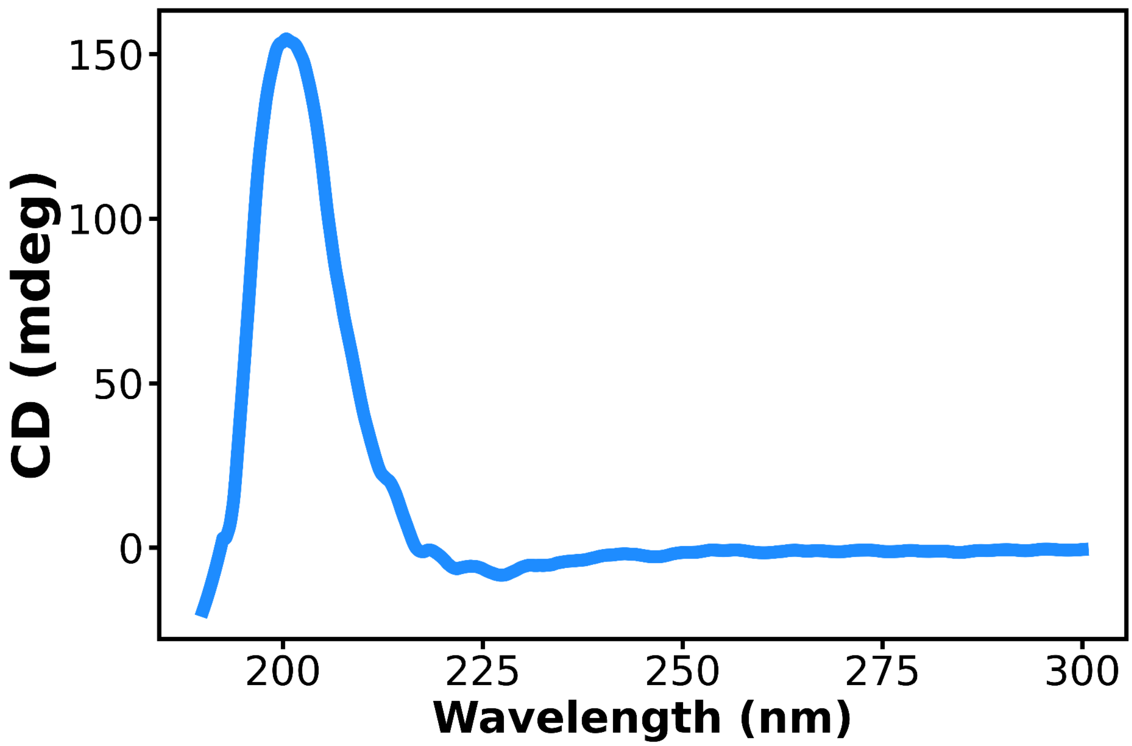


Figure S8: CD spectrum of aged (11-15 days) assembly of Ac-KE. [Ac-KE] = 200 μM.

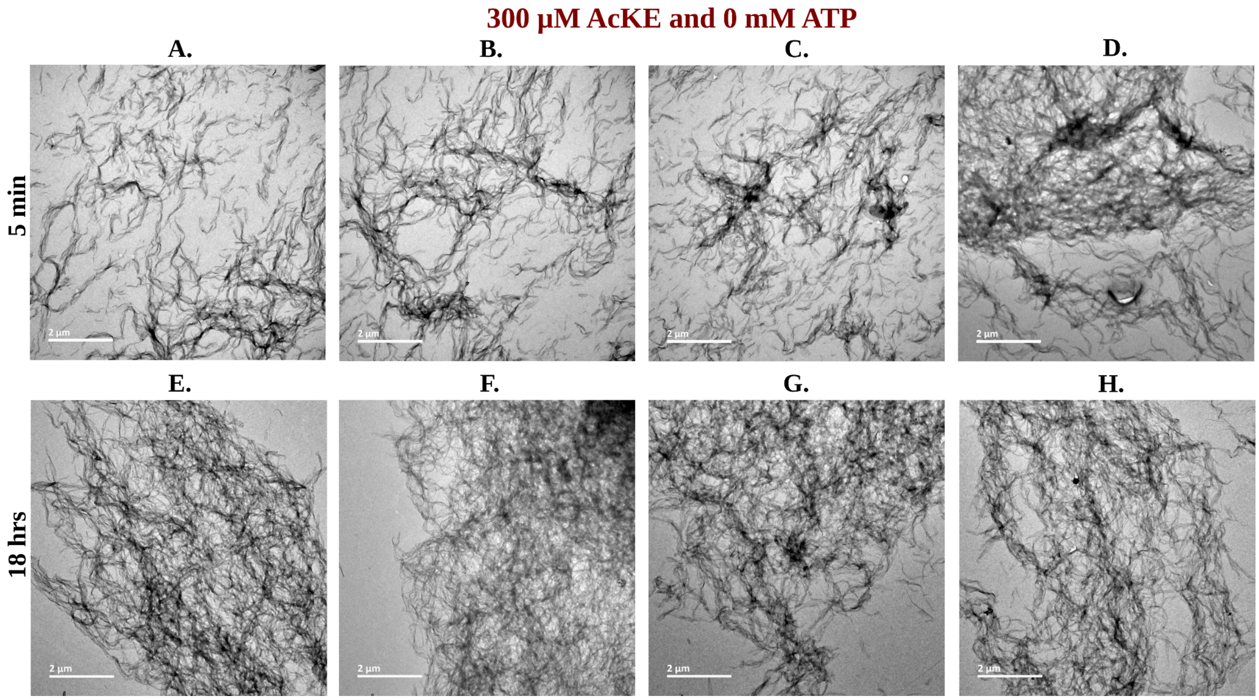


Figure S9: Library of TEM micrographs of 300 µM AcKE in 0 mM ATP at 5 min (figure A-D) and 18 hr (figure E-H).

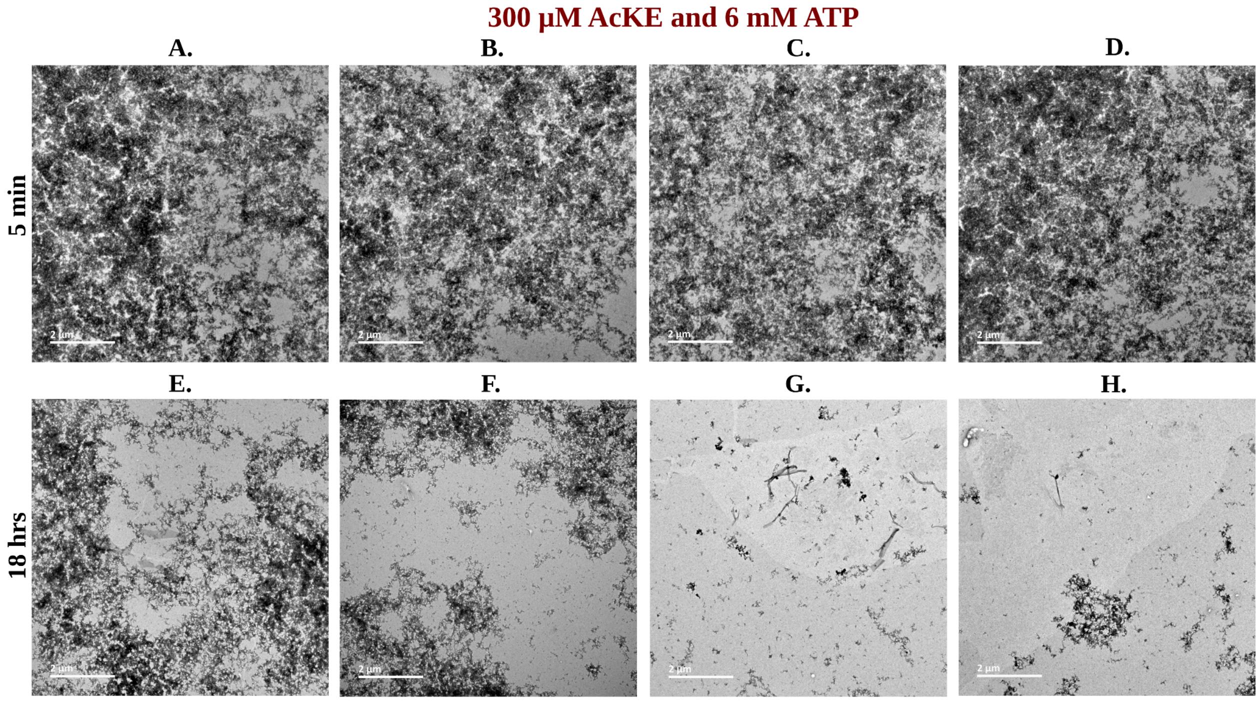


Figure S10: Library of TEM micrographs of 300 µM AcKE in 6 mM ATP at 5 min (figure A-D) and 18 hr (figure E-H).

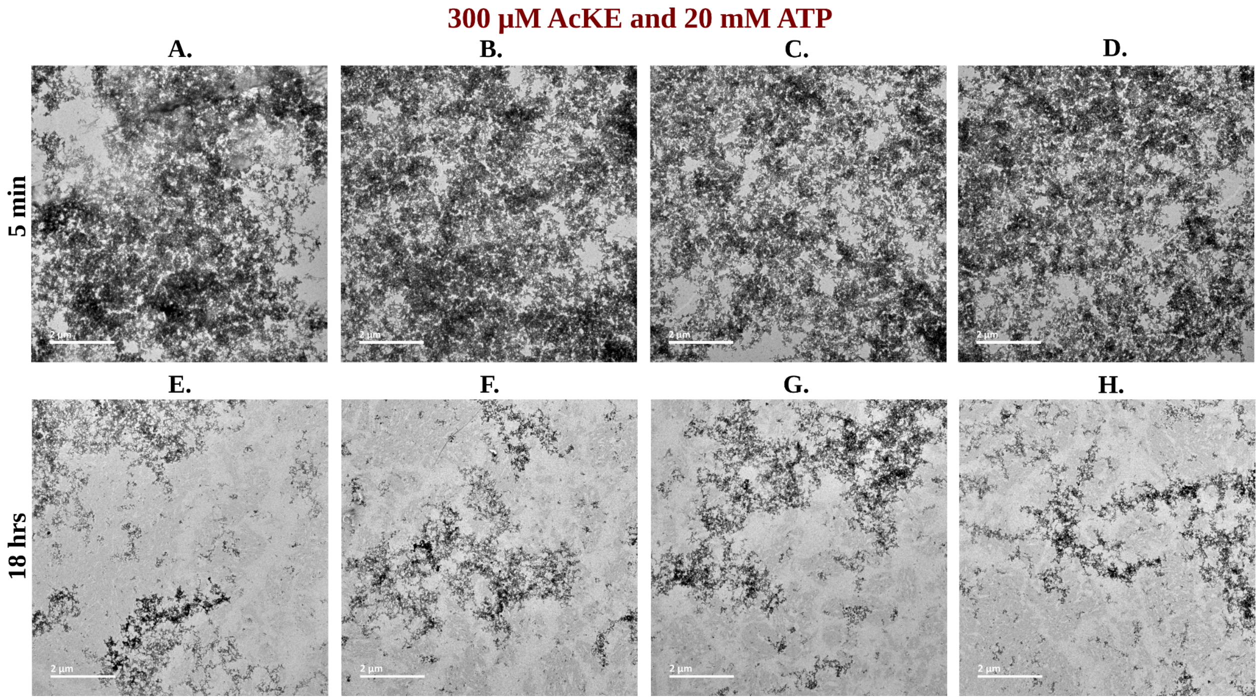


Figure S11: Library of TEM micrographs of 300 µM AcKE in 20 mM ATP at 5 min (figure A-D) and 18 hr (figure E-H).

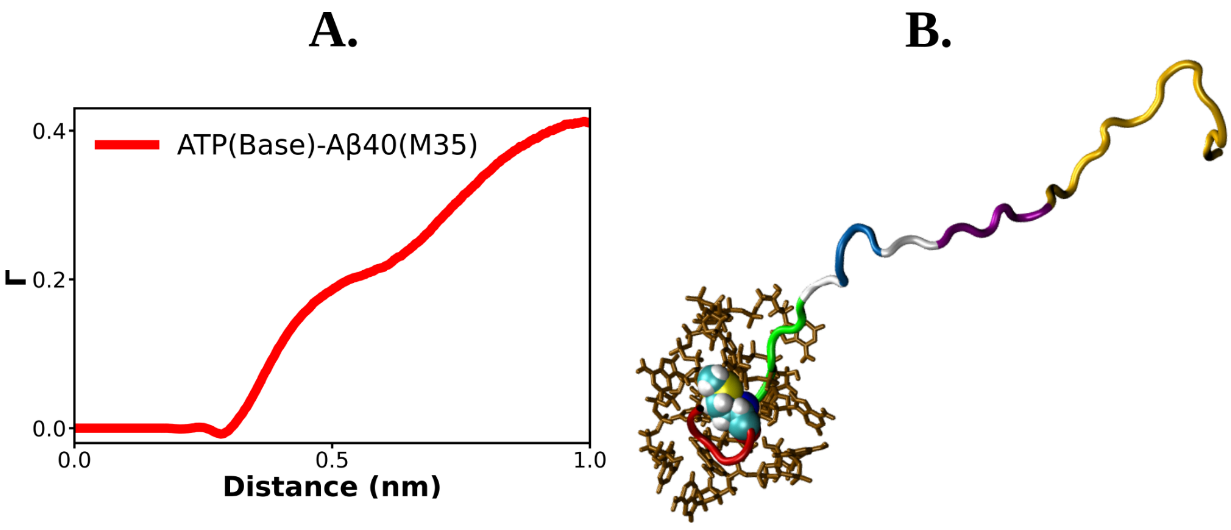


Figure S12: Figure A shows the preferential interaction coefficient of ATP with M35 residue of Aβ40. ATP molecules interacting with M35 residue of Aβ40 is shown with a snapshot in figure B.

**Sequence based analysis:**

**Ubiquitin (pdb id: 1ubq):**
MQIFVKTLT**GKT**ITLEVEPSDTIENVKAKIQDKEGIPPDQQRLIFA**GKQLEDGRT**LSDYNI
QKESTLHLVLRLRGG

**Malate dehydrogenase (pdb id: 4mdh):**
SEPIRVLVTGAAGQIAYSLLYSIGNGSVF**GKDQPIILVLLDIT**PMMGVLDGVLMELQDCA
LPLLKDVIATDKEEIAFKDLDVAILVGSMPRRDGMERKDLLKANVKIFKCQGAALDKYA
KKSVKVIVVGNPANTNCLTASKSAPSIPKENFSCLTRLDHNRAKAQIALKLGVTSDDVKN
VIIWGNHSSTQYPDVNHAKVKLQAKEVGVYEAVKDDSWLKGEFITTVQQRGAAVIKAR
KLSSAMSAAKAICDHVRDIWFGTPEGEFVSMGIISDGNSYGVPDDLLYSFPVTIKDKTWK
IVEGLPINDFSREKMDLTAKELAEEKETAFEFLSSA

**TDP-43 RRM (pdb id: 4bs2):**
**GSHMASKT**SDLIVLGLPWKTTEQDLKEYFSTFGEVLMVQVKKDLKTGHSKGFGFVRFT
EYETQVKVMSQRHMIDGRWCDCKLPNSKQSQDEPLRSRKVFVGRCTEDMTEDELREFF
SQYGDVMDVFIPKPFRAFAFVTFADDQIAQSLCGEDLIIKGISVHISNAEPKHNSNRQ

**Trp-cage (pdb id: 1l2y):**
NLYIQWLKDGGPSSGRPPPS

**Aβ40:**
DAEFRHDSGYEVHHQKLVFFAEDVGSNKGAIIGLMVGGVV
